# A novel cholinergic neural pathway and its role in the drug relapse

**DOI:** 10.1101/2021.05.12.443812

**Authors:** Teng He, Wenwen Chen, Yu Fan, Xing Xu, Zilin Wang, Nanqin Li, Hao Guo, Xue Lu, Feifei Ge, Xiaowei Guan

## Abstract

The lateral parabrachial nucleus (LPB) is critical hub implicated in the control of food intake, reward and aversion. Here, we identified a novel cholinergic projection from choline acetyltransferase (ChAT)-positive neurons in external portion of the lateral parabrachial nucleus (eLPB^ChAT^) to γ–aminobutyric acid (GABA) neurons in central nucleus of amygdala (CeA^GABA^), activation of which could block methamphetamine (METH)-primed conditioned place preference (CPP) in mice.

---

Drug relapse is a big clinic challenge in the treatment of addiction, but its neural circuit mechanism is far from fully understood. The LPB is critical hub implicated in the control of food intake, reward and aversion (*1–3*). A recent study found there seem to exist the ChAT-positive neurons in LPB (*4*), yet their projections and physiological function has not been explored. The central amygdala (CeA) is the main nucleus that receive the outputs from LPB (*5, 6*), and consists of around 95% GABA neurons (CeA^GABA^) (*7*). The neural circuits between LPB and CeA (LPB-CeA) has been implicated in the process of reward and addiction (*1, 8–10*), while most of these studies focus on the glutamatergic projections from calcitonin gene-related peptide (CGRP)-positive neurons in the LPB to CeA (*2, 11, 12*). Here, we found a novel cholinergic projection from LPB to CeA, in which ChAT-positive neurons are mainly located in external portion of LPB (eLPB, Fig. 1a) and more than 50% neurons in eLPB are ChAT-positive neurons (eLPB^ChAT^, Fig. 1b). Cholinergic pathways in the brain, including in the CeA, have been believed to be the classical neurotransmission targets that invoveld in drug-seeking behaviors (*13, 14*). In the study, we dissected the anatomical structure and physiological function of eLPB^ChAT^-CeA pathway, and explored its role in the process of METH relapse.

**Figure 1.**
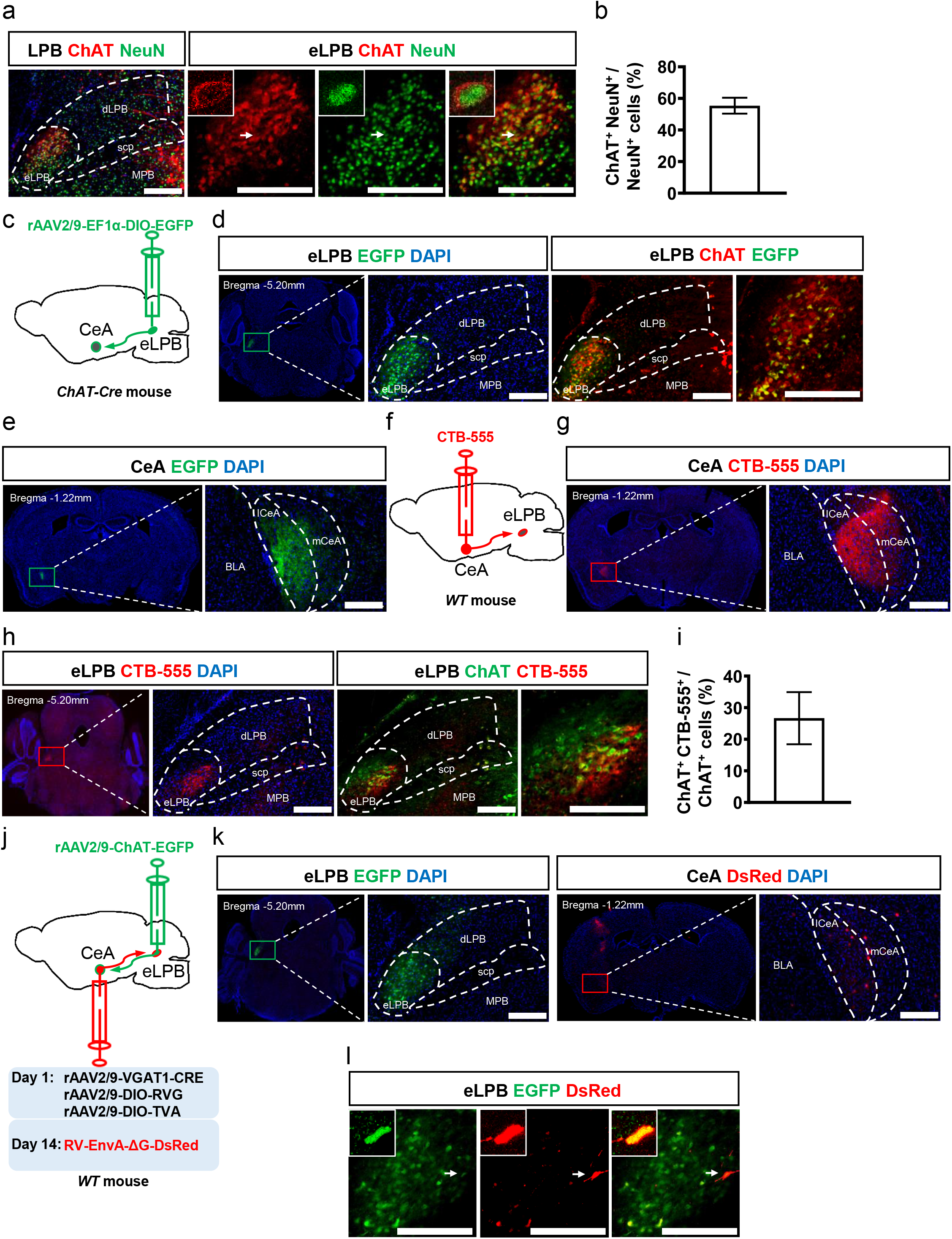
Anatomical dissection of the eLPB ^ChAT^-CeA pathway. **a,** Immunohistochemistry for ChAT/NeuN in the eLPB of WT mice. **b,** The percentage of ChAT-positive cells relative to NeuN-labeled cells in the eLPB. **c,** Schematic of the rAAV2/9-EF1α-DIO-EGFP injection in ChAT-Cre mice. **d**, Representative images of EGFP-labled virus and ChAT-positive neurons in the eLPB of ChAT-Cre mice. **e**, Representative images of EGFP-labeled viral expression within the CeA. **f**, Schematic of CTB-555 injection in WT mice. **g**, Representative images of CTB-555 infusion in the CeA. **h**, CTB-555-transfected and ChAT-positive neurons in the eLPB. **i**, The percentage of CTB-555-transfected cells out of ChAT-positive neurons in the eLPB. **j**, Schematic of viral transfection in the eLPB and CeA of WT mice. **k**, Representative images of rAAV2/9-ChAT-EGFP injection into eLPB and RV-EnvA-ΔG-DsRed injection into CeA. **l**, Representative images of EGFP-labeled and DsRed-labeled viral signals that co-expressed within the eLPB. Scale bar, 400 μm.

To examine the structural projections of eLPB^ChAT^ to CeA^GABA^, we first injected Cre-dependent trans-tracing virus (rAAV2/9-EF1α-DIO-EGFP) into eLPB in ChAT-Cre mice (Fig. 1c, d), and found that eLPB^ChAT^ mainly send projections to lateral region of CeA (lCeA) (Fig. 1e). By stereotaxic injection of retrograde-tracing virus (CTB-555) into CeA in Wild-Type (WT) mice (Fig. 1f, g), we found that around 27% eLPB^ChAT^ neurons are co-labeled with CTB-555 virus from CeA (Fig. 1h, i). In order to confirm the anatomical connection of eLPB^ChAT^ with CeA^GABA^, the anterograde tracing virus specific for ChAT neurons (rAAV2/9-ChAT-EGFP) and the retrograde tracing virus specific for GABA neurons with helper and CRE virus system (rAAV2/9-VGAT1-CRE, rAAV2/9-DIO-RVG, rAAV2/9-DIO-TVA, RV-EnvA-ΔG-DsRed) were injected into eLPB and into CeA in WT mice (Fig. 1j, k). As shown in Fig. 1l, there exist direct projections from eLPB^ChAT^ to CeA^GABA^ (eLPB^ChAT^-CeA^GABA^), as evidenced by the co-transfected ChAT-EGFP virus and retrograde-DsRed virus in eLPB^ChAT^ neurons.

To characterize the functional innervation of eLPB^ChAT^-CeA^GABA^ pathway, the Gq (rAAV2/9-ChAT-hM3D) virus were infused into the bilateral eLPBs to genetic activation of eLPB^ChAT^ neurons (eLPB-Gq) and Go (rAAV2/9-ChAT) virus were used as control (eLPB-Go). In the same mouse, the virus (rAAV2/9-VGAT1-EGFP) were injected into CeA to specifically labeling the CeA^GABA^ neurons (Fig. 2a). After clozapine-N-oxide (CNO) treatment, the frequency of spontaneous action potentials (sAP) of eLPB^ChAT^ neurons were significantly increased when compared to its baseline (Fig. 2b), along with the increased frequency of the spontaneous EPSC (sEPSC) in CeA^GABA^ neurons but which was inhibited by the nicotinic acetylcholine receptors (nAChRs) antagonist mecamylamine (MEC, Fig. 2c), indicating that eLPB^ChAT^ neurons could positively innervate the CeA^GABA^ neurons at least partly by cholinergic projections. To further confirm the above results *in vivo*, fiber photometry of calcium signal was recorded in CeA^GABA^ neurons by injecting rAAV2/9-VGAT1-GCaMp6m into the CeA of WT mice (Fig. 2d, e). As expected, a sustained increase in calcium signals at 30, 35, 40, 45 and 50 min were observed after CNO-activated eLPB^ChAT^ neurons (Fig. 2f, g and h), suggesting the positive innervation of eLPB^ChAT^-CeA^GABA^ pathway.

**Figure 2.**
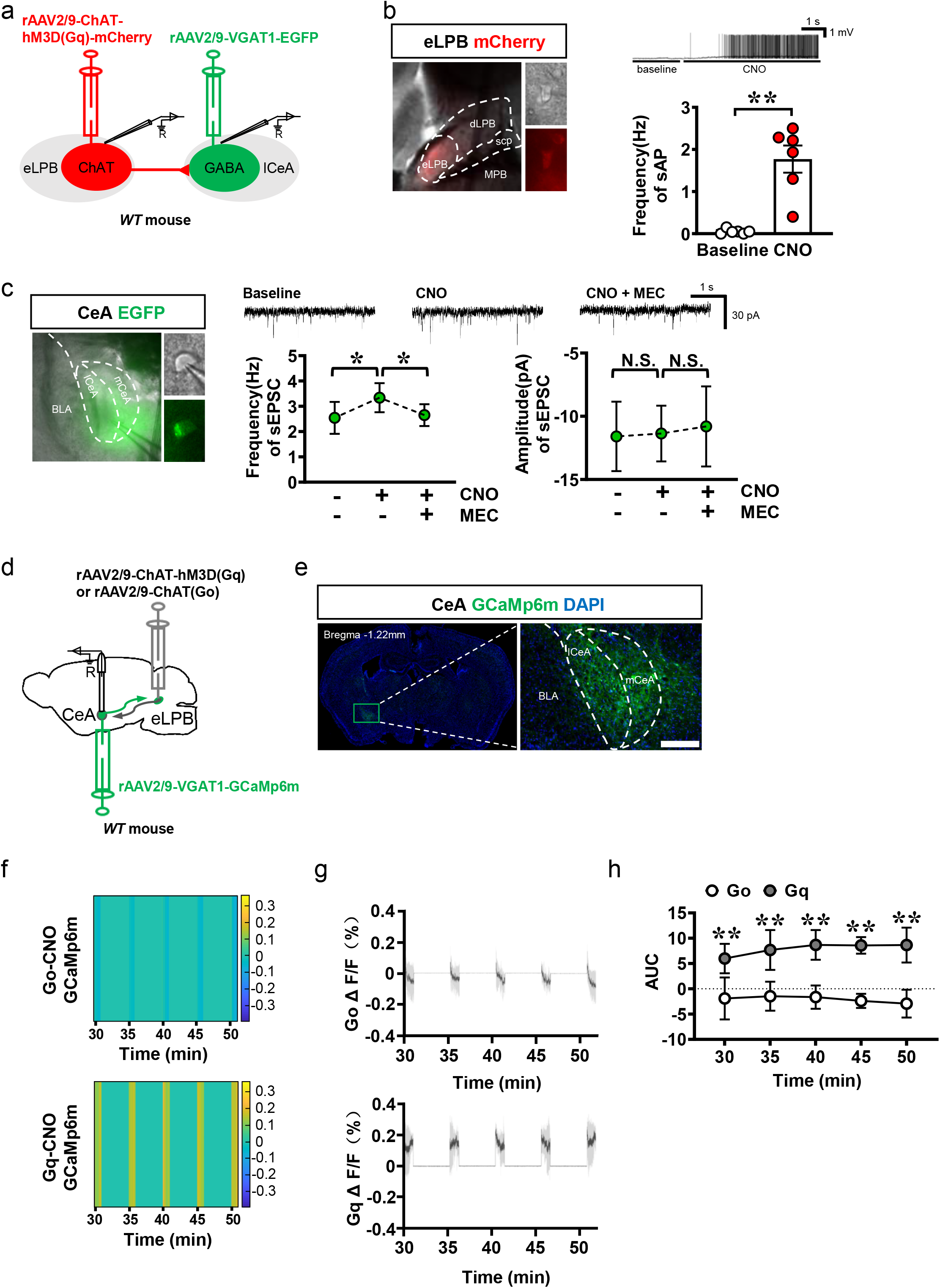
The physiological function of eLPB^ChAT^-CeA^GABA^ pathway. **a**, Schematic of the viral transfection in WT mice and the patch-clamp recording in brain slices. **b**, Representative images of patch-clamp recording on the rAAV2/9-ChAT-hM3D(Gq)-mCherry-transfected neurons in the eLPB (left), sample traces and summarized data of sAP in eLPB^ChAT^ neurons (right) after CNO treatment (n = 6 cells from 6 mice, \*\**P* = 0.0038). **c**, Representative images of patch-clamp recording on the rAAV2/9-VGAT1-EGFP-transfected neurons in the CeA (left), sample traces and summarized data of frequency (middle, n = 6 cells from 6 mice, \**P* = 0.0209 vs. baseline, \**P* = 0.0145 vs. CNO) and amplitude (right, n = 6 cells from 6 mice, ^N.S^.*P* = 0.9756 vs. baseline, ^N.S^.*P* = 0.7498 vs. CNO) of sEPSC in response to CNO and MEC. **d**, Schematic of viral transfection and optical fiber implantation in the eLPB and CeA. **e**, Representative images of rAAV2/9-VGAT1-GCaMp6m-transfected neurons in the CeA. **f**, Heatmap of GCaMp6m fluorescence at 30, 35, 40, 45 and 50min after CNO administration. **g**, Quantification of the peak ΔF/F. **h**, Go and Gi-evoked AUC (n = 6, F _(4, 16)_=1.289, \*\**P*_(30)_ = 0.0078, \*\**P*_(35)_ = 0.0044, \*\**P*_(40)_ = 0.0034, \*\**P*_(45)_ = 0.0021, \*\*\**P*_(50)_ = 0.0005 vs. Go). Scale bar, 400 μm.

In order to detect the role of eLPB^ChAT^ neurons and its projections to CeA in the process of METH relapse, METH-primed CPP were set up in mice (Fig. 3a, b). To assess the role of eLPB^ChAT^ neurons in METH-primed reinstatement of CPP, we used eLPB-Gq and eLPB-Go mice models (Fig. 3c, d). As shown in Fig. 3e-f, a single challenge of METH successfully reinstated the METH CPP in eLPB-Gq mice, but failed to reinstate that in eLPB-Go mice after systemic administration of CNO treatment. To rule out the role of eLPB^ChAT^-CeA pathway in METH-primed CPP, we employed a combinational viral strategy by infusion of rAAV2/retro-Cre-EGFP into CeA and infusion of Cre-dependent rAAV2/9-ChAT-DIO-hM3D(Gq)-mCherry into eLPB (Fig. 3g, h). As shown in Fig. 3i-j, chemico-genetic activation of eLPB^ChAT^ terminals-projecting into the CeA by CNO obviously block the METH-primed CPP behaviors in mice. These findings suggest that eLPB^ChAT^ neurons and its projections to CeA control the METH relapse behaviors.

**Figure 3.**
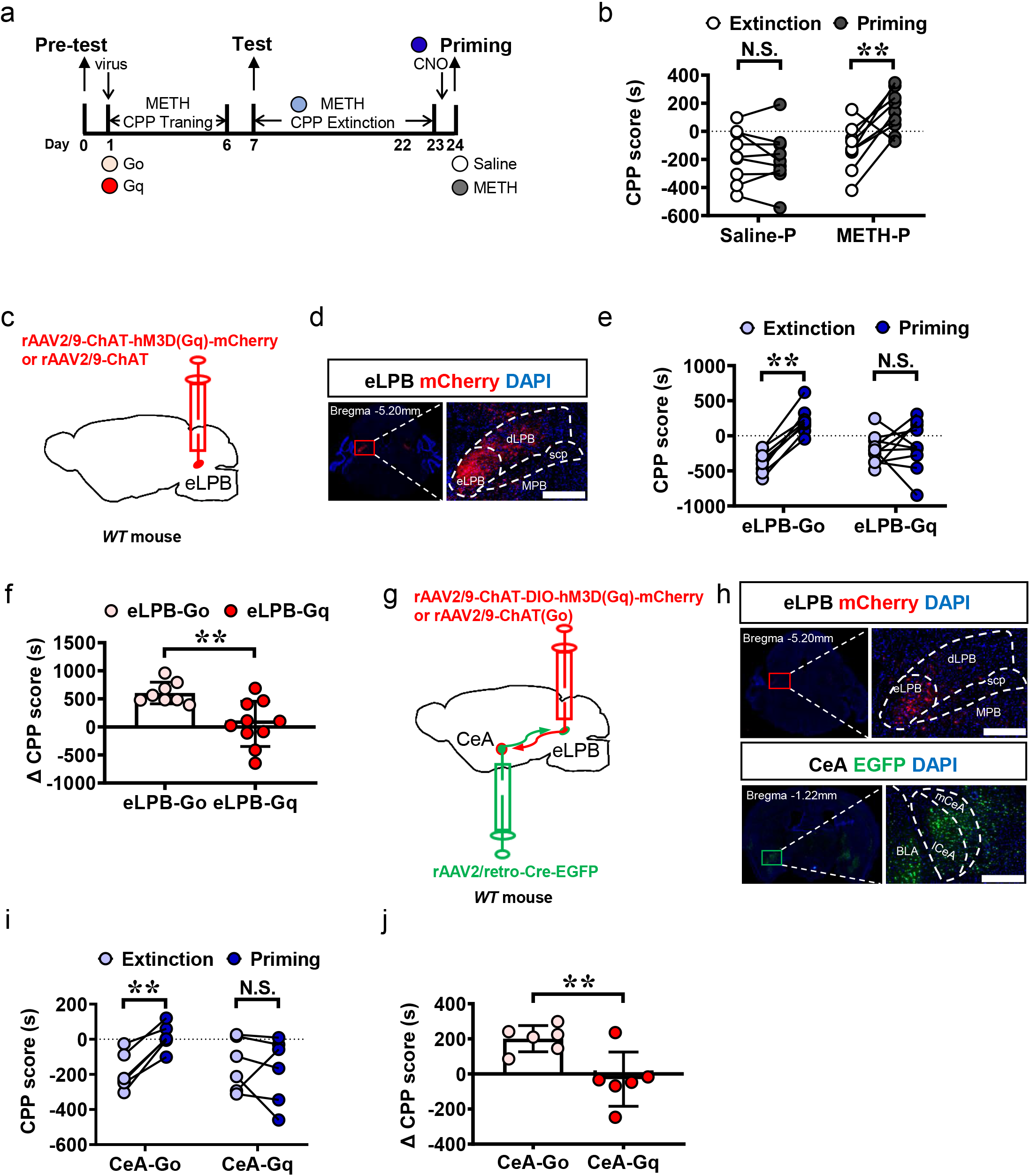
Effects of chemico-genetic activating eLPB^ChAT^-CeA pathway on METH-primed CPP. **a**, Experimental design and timeline. **b**, CPP scores in METH-treated WT mice. (Saline-primed METH CPP, n = 9, ^N.S^.*P* = 0.5216 vs. Extinction; METH-primed METH CPP, n = 9, \*\**p* = 0.006 vs. Extinction). **c**, Schematic of the viral transfection of WT mice. **d**, Representative images of rAAV2/9-ChAT-hM3D(Gq)-mCherry or rAAV2/9-ChAT(Go) injection in the eLPB. **e & f**, The CPP and ΔCPP scores after activating eLPB^ChAT^ neurons during METH-primed CPP (eLPB-Go group, n = 8, \*\**P* <0.0001 vs. Extinction; eLPB-Gq group, n = 10, ^N.S^.*P* = 0.6808 vs. Extinction; n = 18, \*\**P* = 0.0027 vs. Go). **g**, Schematic of the viral transfection in the eLPB and CeA of WT mice. **h**, Representative images of rAAV2/9-ChAT-hM3D(Gq)-mCherry or rAAV2/9-ChAT(Go) injection in the eLPB (top), and rAAV2/retro-Cre-EGFP within the CeA (bottom). **i & j**, The CPP and ΔCPP scores after activating eLPB^ChAT^-CeA pathway during METH-primed CPP. (CeA-Go group, n = 6, \*\**P* = 0.0012 vs. Extinction; CeA-Gq group, n = 6, ^N.S^.*P* = 0.6573 vs. Extinction; n = 12, \*\**P* = 0.0082 vs. Go). Scale bar, 400 μm.

In summary, we found a novel cholinergic pathway from LPB^ChAT^ neurons to CeA^GABA^ neurons. Activation of eLPB^ChAT^-CeA pathway benefits for the treatment of METH relapse. The key molecules of the eLPB^ChAT^-CeA^GABA^ pathway underlying drug relapse should be studied in the future study.

## Methods

### Animals

C57BL/6 WT and ChAT-Cre male mice weighing 25-35g were used. Animals were housed at constant humidity (40~60%) and temperature (24 ± 2 °C) with a 12-hour light/dark cycle and allowed free access to food and water. All procedures were carried out in accordance with the National Institutes of Health Guide for the Care and Use of Laboratory Animals and approved by the Institutional Animal Care and Use Committee (IACUC) at Nanjing University of Chinese Medicine.

### Virus injection

The mice were fixed in a stereotactic frame (RWD, Shenzhen, China) under 2% isoflurane anesthesia. A heating pad was used to maintain the core body temperature of the mice at 37 °C. Unless otherwise noted, a volume of 100 nl virus was injected using calibrated glass microelectrodes connected to an infusion pump (Drummond, Alabama, USA) at a rate of 20 nl/min. The coordinates were defined as dorsoventral (DV) from the brain surface, anterior-posterior (AP) from bregma, and mediolateral (ML) from the midline (in mm). The stereotaxic coordinates of eLPB are as following: AP, −5.20 mm, ML, ±1.55 mm and DV, −3.60 mm. The stereotaxic coordinates of CeA are as following: AP, −0.90 mm; ML, ±2.70 mm; DV, −4.55 mm. All viruses were packaged by BrainVTA (Wuhan, China).

For anterograde tracing, rAAV2/9-EF1α-DIO-EGFP-WPRE-hGHpA (rAAV2/9-EF1α-DIO-EGFP, 2.04E+12vg/ml) were injected into the eLPB of ChAT-Cre mice, which enabled the Cre enzyme to spread anterogradely and monosynaptically into the soma of eLPB neurons. The expression of EGFP was detected in the CeA, and co-staining with ChAT antibody in the LPB after 3-week transfection. In the patch-clamp recording experiment, the rAAV2/9-ChAT-hM3D(Gq)-mCherry-WPREs (rAAV2/9-ChAT-hM3D(Gq)-mCherry, 5.54E+12vg/ml) was injected into eLPBs, and rAAV2/9-VGAT1-EGFP-WPRE-hGHpA (rAAV2/9-VGAT1-EGFP, 5.31E+12vg/ml) was delivered into the CeAs of WT mice. We used 2 cohorts of WT mice in the CPP experiment. In the cohort-1 mice, the rAAV2/9-ChAT-hM3D(Gq)-mCherry-WPREs (rAAV2/9-ChAT-hM3D(Gq)-mCherry, 5.54E+12vg/ml) or rAAV2/9-ChAT-WPRE-hGHpA (rAAV2/9-ChAT(Go), 5.50E+12vg/ml) was injected into eLPB. In cohort-2 mice, the Cre-dependent virus rAAV2/9-ChAT-DIO-hM3D(Gq)-mcherry-WPREs (rAAV2/9-ChAT-DIO-hM3D(Gq)-mcherry, 5.08E+12vg/ml) or rAAV2/9-ChAT-WPRE-hGHpA (rAAV2/9-ChAT(Go), 5.50E+12vg/ml) was delivered into the eLPBs and rAAV2/retro-hSyn-CRE-EGFP-WPRE-hGHpA (rAAV2/retro-Cre-EGFP, 5.25E+12vg/ml, 150 nL) was co-injected into the CeAs.

For retrograde tracing, the retrograde tracer CTB-555 (1 μg/μl) was used to trace the eLPB^ChAT^ neurons projecting to the CeA. In another cohort of WT mice, helper viruses that contained rAAV2/9-VGAT1-CRE-WPRE-pA (rAAV2/9-VGAT1-CRE, 3.63E+12vg/ml, 25 nl), rAAV2/9-EF1α-DIO-H2B-EGFP-T2A-TVA-WPRE-hGHpA (rAAV2/9-DIO-TVA, 5.56E+12vg/ml, 25 nl) and rAAV2/9-EF1α-DIO-oRVG-WPRE-hGHpA (rAAV2/9-DIO-RVG, 5.29E+12vg/ml, 50 nl) were co-injected into the CeA of WT mouse. The rAAV2/9-ChAT-EGFP-WPRE-pA (rAAV2/9-ChAT-EGFP, 2.15E+12vg/ml) was delivered into the eLPB in the same mice. After 2 weeks, the rabies virus RV-EnvA-ΔG-DsRed (2.00E+08 IFU/ml) was injected into the same site in the CeA. The modified rabies virus cannot infect neurons and spread retrogradely without the TVA and rabies glycoprotein component, which is provided by AAV helpers.

All mice were perfused with 0.9% saline, followed by 4% PFA. Images of the fluorescence signals were visualized using a Leica DM6B upright digital research microscope (Leica, Germany) or Leica TCS SP8 STED (Leica, Germany). Animals with missed injections were excluded in the present study.

### Immunohistochemistry

The mice were deeply anesthetized with 10% chloral hydrate (0.2 ml, i.p.) and sequentially perfused with 0.9% saline and 4% PFA. The brains were removed and post-fixed in 4% PFA at 4 °C overnight. After cryoprotection of the brains with 30% (w/v) sucrose, coronal sections (30 μm) were cut on a cryostat (Leica, Germany) and used for immunofluorescence. The sections were incubated in 3% (v/v) Triton X-100 for 0.5 h, blocked with donkey serum for 1.5 h at room temperature, and incubated with the following primary antibodies: goat anti-ChAT (1:200, Millipore, USA), rabbit anti-NeuN (1:200, Cell Signaling Technology, USA) at 4 °C for 24 h, followed by the corresponding fluorophore-conjugated secondary antibodies for 2 h at room temperature. The following secondary antibodies were used here: Alexa Fluor 555-labeled donkey anti-goat secondary antibody (1:500, Invitrogen, USA), Alexa Fluor 488-labeled donkey anti-goat secondary antibody (1:500, Invitrogen, USA), Alexa Fluor 488-labeled donkey anti-rabbit secondary antibody (1:500, Invitrogen, USA). Fluorescence signals were visualized using a Leica DM6B upright digital research microscope (Leica, Germany) or Leica TCS SP8 STED (Leica, Germany).

### Brain slice electrophysiology

Mice were anesthetized with 2% isoflurane and perfused with ice-cold oxygenated modified NMDG artificial cerebrospinal fluid (NMDG ACSF) that contained (in mM) 92 N-methyl-d-glucamine (NMDG), 2.5 KCl, 1.2 NaH_2_PO_4_·2H_2_O, 30 NaHCO_3_, 20 HEPES, 25 Glucose, 5 Sodium ascorbate, 2 Thiourea, 3 Sodium Pyruvate, 10 MgSO_4_, 0.5 CaCl_2_·2H_2_O (the pH was adjusted to 7.3-7.4 with HCl and saturated with 95% O_2_/5% CO_2_). Coronal slices (200 μm) that contained the LPB or CeA were sectioned on a vibrating microtome (VT1000s, Leica, Germany) in ice-cold cutting solution. The brain slices were initially incubated in NMDG ACSF for 9 min at 37 °C, followed by N-2-hydroxyethylpiperazine-N-2-ethanesulfonic acid (HEPES) ACSF that contained (in mM) 86 NaCl, 2.5 KCl, 1.2 NaH_2_PO_4_·2H_2_O, 35 NaHCO_3_, 20 HEPES, 25 Glucose, 5 Sodium ascorbate, 2 Thiourea, 33 Sodium Pyruvate, 1 MgSO_4_, 2 CaCl_2_·2H_2_O (the pH was adjusted to 7.3-7.4 with HCl and saturated with 95% O_2_/5% CO_2_) for at least 30 min at 37 °C and then at room temperature for 1 h. During recordings, brain slice was continuously perfused with oxygenated ACSF that contained (in mM) 119 NaCl, 2.5 KCl, 1 NaH_2_PO_4_·2H_2_O, 26.2 NaHCO_3_, 1.3 MgCl_2_, 2.5 CaCl_2_, 11 D-Glucose (the pH was adjusted to 7.3-7.4 with HCl and osmolarity was 290-295 mOsm). The slices were recorded while they were bath-perfused (2.5 ml/min) at 30°C–32°C by an in-line solution heater (TC-324C, Warner Instruments, USA) in the same ACSF solution with the addition of 50 μM picrotoxin to block GABAA receptors. Internal solution (in mM) 130 CsMeSO_4_, 10 NaCl, 10 EGTA, 4 Mg-ATP, 0.3 Na-GTP, 10 HEPES (the pH was adjusted to 7.3-7.4 with HCl) was used for spontaneous excitatory postsynaptic current (sEPSC) recordings.

The whole-cell voltage-clamp recordings (Vholding= −70mV) were carried out in mCherry-positive neurons in the eLPB (red) or EGFP-positive neurons in the CeA (green) using an Olympus BX51WIF microscope optical system combined with a Digital Camera (C11440-42U, Hamamatsu, Japan). During sAP recording in the eLPB, CNO (10 μM) was used in the recording ACSF solution. During sEPSC recording in the CeA, CNO (10 μM) and MEC (5 μM) were used in the recording ACSF solution. Patch pipettes (3-5 MΩ) were pulled from borosilicate glass capillaries (Sutter Instruments, USA) with an outer diameter of 1.5 mm on a four-stage horizontal puller (P1000, Sutter Instruments, USA). The signals were acquired with a Multiclamp 700B amplifier (Molecular Devices, USA), low-pass filtered at 4.0 kHz, digitized at 10 kHz and analyzed with Clampfit 10.5 software (Molecular Devices, USA).

### Calcium fiber photometry

For the photometry recording, Cre-dependent AAV vectors carrying fluorescent calcium indicator rAAV2/9-VGAT1-GCaMp6m-WPRE-hGHpA (rAAV2/9-VGAT1-GCaMp6m, 5.25E+12vg/ml) was injected into CeAs, and the rAAV2/9-ChAT-hM3D(Gq) or rAAV2/9-ChAT(Go) was injected into eLPBs. An optical fiber (200μm outer diameter, 0.37 numerical aperture, Inper Ltd., China) was placed 150 μm above the viral injection site. Calcium-dependent fluorescence signals were obtained by stimulating cells expressing GCaMp6m with a 470 nm LED (35~40 μW at fiber tip) while calcium-independent signals were obtained by stimulating these cells with a 405 nm LED (15~20 μW at fiber tip). The LED lights of 470 nm and 405 nm were alternated at 66 fps and light emission was recorded using sCMOS Camera containing the entire fiber bundle (2 m in length, NA = 0.37, 200 μm core, Inper Ltd.). The analog voltage signals fluorescent was filtered at 30 Hz and digitalized at 100 Hz. The calcium-dependent fluorescence signals were recorded by Inper Studio Multi Color EVAL15 software (Inper Ltd., China) at 30min, 35min, 40min, 45min and 50min after CNO (2mg/kg, i.p.) treatment with 1min recording duration and 4min interval of each trial, and the raw data were analyzed using Inper Data Process V0.5.9 (Inper Ltd., China).

The values of ΔF/F are calculated by (F-F0)/F0, where F0 is the baseline fluorescence signal and F is the real-time fluorescence signal at 30min, 35min, 40min, 45min and 50min after CNO treatment. The area under curve (AUC) is the area value under recording duration related to corresponding baseline at every trial.

### CPP

METH CPP procedures were performed in the TopScan3D CPP apparatus (CleverSys, VA, USA), which is constructed of two distinct chambers (30.5 × 25 × 32 cm each) separated by a removable guillotine door. The CPP procedure consisted of five phases: the pre-conditioning test (Pre-test, day 0), conditioning (CPP training, days 1-6), post-conditioning test (Test, day 7), extinction training (Extinction, day 8-23), and reinstatement test (Priming, day 24). During CPP training conditioning, mice was injected with METH (3 mg/kg, i.p.) once daily. After each injection, the mice were confined to one conditioning chamber (drug-paired chamber) for 45 min and then returned to their home cages. To train CPP extinction, mice were confined to the drug-paired chamber for 45 min once daily without any drug treatment. During the Pre-test, Test, and Priming, mice were allowed to freely access two chambers for 15 min. The CPP score is the time spent in the drug-paired chamber, and the ΔCPP score is the value of the time spent in drug-paired chamber minus that in non-drug-paired chamber.

### Statistical Analysis

Statistical analysis was carried out using GraphPad Prism 8.0 software. All data are presented as the mean ± SD (standard deviation). The data of Aps, sEPSC and CPP scores were analyzed with paired t-tests. The data of calcium signals were analyzed by repeated measures of two-way analysis of variance with Tukey post hoc tests. The data of ΔCPP scores were analyzed by unpaired t-tests. Statistical significance was set as *P* < 0.05.

## Acknowledgements

This work is supported by National Natural Science Foundation of China (No. 82071495 and No. 81901353), the Priority Academic Program Development of Jiangsu Higher Education Institutions (Integration of Chinese and Western Medicine), and Natural Science Foundation of Jiangsu Province, China (No. BK20201398 and No. BK20190805). We thank Prof. Minmin Luo lab, National Institute of Biological Science, Beijing, China, for giving ChAT-Cre mice used in the study.

## Author contributions

Guan X and Ge F developed the overall concept and supervised the experiments. Guan X, Ge F, Fan Y and He T co-analyzed data and co-wrote the manuscript. He T, Chen W, Fan Y performed virus-injected mice models, morphological experiments, fiber photometry and behavioral tests. Xu X and Wang Z performed patch-clamp electrophysiology. Li N, Guo H and Lu X helped with behavioral, morphological and virus studies.

